# A Bayesian Zero-Inflated Binomial Regression and Its Application in Dose-Finding Study

**DOI:** 10.1101/676809

**Authors:** Puntipa Wanitjirattikal, Chenyang Shi

**Affiliations:** King Mongkut’sInstitute of Technology Ladkrabang, Thailand; Celgene Corporation, USA

**Author notes:** These two authors contributed equally to this work.

**Keywords:** dose-limiting toxicity, maximum-tolerated dose, metropolis algorithm, zero-inflated binomial regression

## Abstract

In early phase clinical trial, finding maximum-tolerated dose (MTD) is a very important goal. Many researches show that finding a correct MTD can improve drug efficacy and safety significantly. Usually, dose-finding trials start from very low doses, so in many cases, more than 50% patients or cohorts do not have dose-limiting toxicity (DLT), but DLT may occur suddenly and increase fast along with just two or three doses. Although some fantastic models were built to find MTD, little consideration was given to those ‘0 DLTs’ and the ‘jump’ of DLTs. We developed a Bayesian zero-inflated binomial regression for dose-finding study based on Hall (2000), which analyses dose-finding data from two aspects: 1) observation of only zeros, 2) number of DLTs based on binomial distribution, so it can help us analyse if the cohorts without DLT have potential possibility to have DLT and fit the ‘jump’ of DLTs.

## 1. Introduction

In clinical trial, finding maximum-tolerated dose (MTD) is one of the chief goals in phase 1 or 2. MTD is generally defined as maximum dose can be tolerated by patients, and the tolerance is usually measured via the probability of dose-limiting toxicity (DLT) which is the toxicity occurred in patients. For example, we have 8 dose levels for a drug, 1 mg, 2, mg, 4 mg, 8 mg, 12 mg, 16 mg, 22 mg, and 35 mg. The first 5 cohorts were enrolled with 3 patients for each, and the last 3 cohorts were enrolled with 6 patients for each. Our data is presented as follows:

|  |  |  |  |  |  |  |  |  |
| --- | --- | --- | --- | --- | --- | --- | --- | --- |
| Doses (mg) | 1 | 2 | 4 | 8 | 12 | 16 | 22 | 35 |
| Number of Patients | 3 | 3 | 3 | 3 | 3 | 6 | 6 | 6 |
| Number of DLTs | 0 | 0 | 0 | 0 | 0 | 1 | 2 | 4 |

From cohort 1 to 5, no DLT occurred, because dose-finding studies usually start from very low doses. 1 DLT occurred at dose level 6, 2 DLTs occurred at dose level 7, and 4 DLTs occurred at dose level 8. More than 50% cohorts in our data do not have DLT. If we define MTD as the dose with 33% of probability of DLT (P(DLT)), then based on the observed P(DLT), 22 mg may be MTD in our study. However, our analysis should be based on the potential P(DLT) curve with prior information (i.e., historical studies) instead of observed curve, because in early phase studies, especially, oncology studies, sample size is always small. O’Quigley (1990) proposed a continual reassessment method (CRM) for MTD finding. This is a very influential method in clinical trial and some basic theories were stated in his paper. The potential P(DLT) curve was assumed to be monotonic with dose levels, and a Bayesian binomial framework was built so that prior information can be incorporated. A significant development of CRM is a two-parameter Bayesian logistic regression proposed by Neuenschwander (2008), which is widely used in pharmaceutical industry. This is a very flexible model for adaptive dose-find design, and covariates can be added in easily (Bailey, 2009). Another logistic based Bayesian model is proposed by Tighiouart (2005). Apparently, binomial regression is the most suitable for DLT-based dose-finding studies, since DLT is a yes/no variable. But so far, to our knowledge, little work has been done to discuss those ‘0 DLTs’ in dose-finding data. Since dose-finding trials usually start from very low doses, more than 50% cohorts or patients may have no DLT, but DLT may occur suddenly and increase fast along with just two or three doses, like our example above. This implies that in this kind of studies, P(DLT) may be fit in two curves, one curve is for 0 DLTs, and the other curve is for non-0 DLTs, and these two curves are not independent. To explore this question, a zero-inflated binomial (ZIB) regression may be a good lever.

ZIB regression is a statistical model to fit binary data with excessive zeros, which was inspired by zero-inflated Poisson regression (Lambert, 1992) and first proposed by Hall (2000). ZIB is a mixture of observation of only zeros and a weighted binomial distribution. Two unknown parameters in ZIB are probability of observation from only zeros and probability of success in binomial distribution, and for regression, logit link functions can be imposed on these two parameters to incorporate covariates. An EM algorithm is given in Hall (2000) for parameter estimation. However, as we introduced before, prior information is very important in dose-finding studies, since our analysis will be based on potential P(DLT) curves with historical information. To incorporate prior information and calculate probabilities of under dose, target dose, and over dose for safety control, it is necessary to develop a Bayesian algorithm for ZIB regression.

In this paper, we developed a Bayesian ZIB (BZIB) regression for dose-finding study based on Hall (2000). In Section 2, we introduced a general Bayesian framework for ZIB regression, and simulations were conducted to evaluate the performance of our Bayesian algorithm. In Section 3, we conducted simulations to assess the accuracy of BZIB regression in dose-finding study, and applied BZIB regression to our data in introduction. Our conclusion is in Section 4.

## 2. BZIB Regression

### 2.1 Bayesian Inference for ZIB regression

First, let us discuss a BZIB regression in a general situation. Assuming we have *N* samples. Let *n*_*i*_ denote the *i*th sample size, and *y*_*i*_ denote the number of successful events of *i*th sample, *i* = 1, 2, …, *N*. ZIB can be written as:

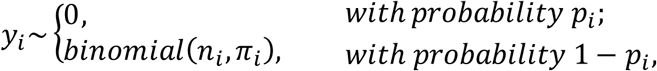

where, *π*_*i*_ is the probability of success in *i*th sample, and *p*_*i*_ is the probability that *y*_*i*_ is from the observation of only zeros. This implies that ZIB regression can be written as:

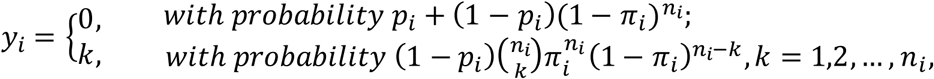

logit links can be imposed on *p*_*i*_ and *π*_*i*_, so *logit*(*p*_*i*_) = ***Z***_*i*_***γ***, and *logit*(*π*_*i*_) = ***X***_*i*_***β***.***Z*** and ***X*** are covariate matrices. Let *u*_*i*_ = 1 when *y*_*i*_ = 0, and *u*_*i*_ = 0 when *y*_*i*_ = 1, the joint density of ZIB regression is:

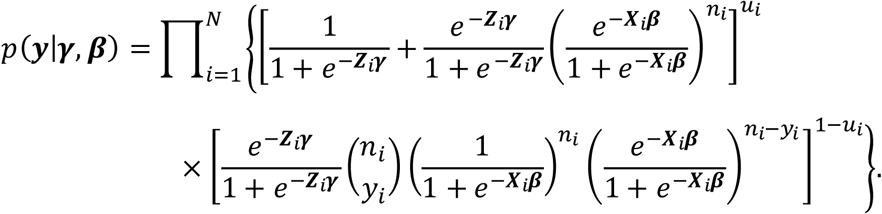

Let *p*(***γ***) and *p*(***β***) denote the prior distribution of ***γ*** and ***β***, respectively. The posterior distribution of ZIB regression is:

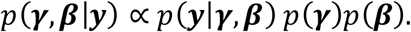

To estimate ***γ*** and ***β***, an easy way is to use metropolis algorithm which is a Markov chain Monte Carlo (MCMC) sampling method (Ghosh, 2006; Metropolis, 1953; Hoff, 2009). Metropolis algorithm requires posterior distributions and candidate parameters from proposal distributions. For simplicity, we usually assume that ***γ*** and ***β*** are independent. We assign normal priors to our parameters, and adopt normal distributions as our proposal distributions, since the range of our parameters are (−∞, +∞). Using un-bold *γ* anD *β* to represent each single parameter in ***γ*** and ***β***, our algorithm is shown as follows:

#### Algorithm

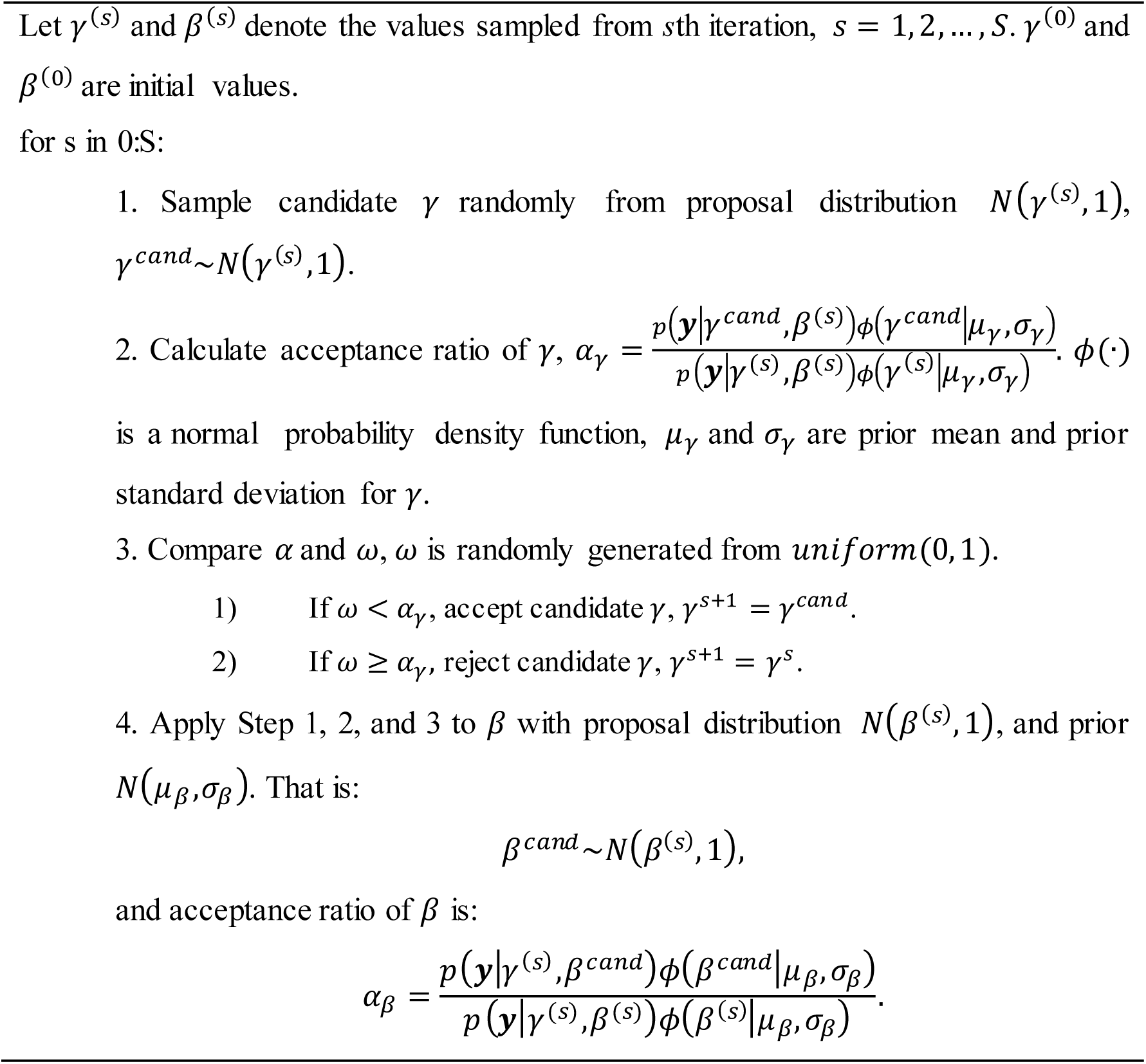

In Step 2, to avoid the overflow of extreme large values and improve the efficiency of computation, we can do log transform for *α*, and the log of the density of ZIB regression is:

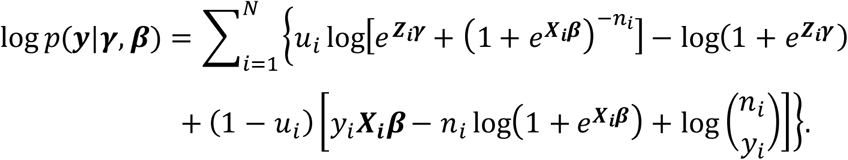

Correspondingly, we compare log *ω* with log *α* in Step 3.

### 2.2 Simulation Study

To assess the performance of Metropolis algorithm for ZIB regression, and the accuracy of our estimation, we conducted 500 simulations on the data generated from ZIB regression with *logit*(*p*_*i*_) = *γ*_0_ + *γ*_1_*X*_*i*_ and *logit*(*π*_*i*_) = *β*_0_ + *β*_1_ *X*_*i*_, and *X*∼*Piosson*(10)/5. A non-informatively normal prior, *N*(0, 10000), was assigned to each *γ*_0_, *γ*_1_, *β*_0_, and *β*_1_. We proposed three cases: 1) *γ*_0_ = 2, *γ*_1_ = −1, *β*_0_ = −4, *β*_1_ = 2; 2) *γ*_0_ = 1, *γ*_1_ = −0.5, *β*_0_ = −1, *β*_1_ = 0.5; 3) *γ*_0_ = 2, *γ*_1_ = −1.5, *β*_0_ = −1.5, *β*_1_ = 1, with sample size of *n* = 100 and *n* = 200. For each simulation, we ran 10000 MCMC iterations with 5000 burn-ins in R 3.4.3. The performance of our algorithm is evaluated by mean and standard deviation (SD) of the estimates from simulations, percentage of bias between true values and estimated values, and coverage probability (CP). Our simulation results are presented in Table 1.

**Table 1:** Simulation Results of BZIB Regression

| Case | n | Parameter | Mean (SD) | Bias (%) | CP |
| --- | --- | --- | --- | --- | --- |
| 1 | 100 | $\gamma_0$ | 2.057 (1.11) | -0.029 | 0.952 |
| | | $\gamma_1$ | -1.043 (0.512) | -0.043 | 0.954 |
| | | $\beta_0$ | -4.131 (0.792) | -0.033 | 0.922 |
| | | $\beta_1$ | 2.062 (0.357) | -0.031 | 0.938 |
| | 200 | $\gamma_0$ | 2.066 (0.752) | -0.033 | 0.912 |
| | | $\gamma_1$ | -1.036 (0.342) | -0.036 | 0.918 |
| | | $\beta_0$ | -4.098 (0.526) | -0.025 | 0.92 |
| | | $\beta_1$ | 2.046 (0.237) | -0.023 | 0.914 |
| 2 | 100 | $\gamma_0$ | 1.1 (0.824) | -0.1 | 0.928 |
| | | $\gamma_1$ | -0.553 (0.386) | -0.106 | 0.932 |
| | | $\beta_0$ | -1.043 (0.526) | -0.043 | 0.936 |
| | | $\beta_1$ | 0.513 (0.242) | -0.026 | 0.914 |
| | 200 | $\gamma_0$ | 1.012 (0.54) | -0.012 | 0.954 |
| | | $\gamma_1$ | -0.509 (0.257) | -0.018 | 0.956 |
| | | $\beta_0$ | -1.024 (0.341) | -0.024 | 0.92 |
| | | $\beta_1$ | 0.513 (0.154) | -0.026 | 0.932 |
| 3 | 100 | $\gamma_0$ | 2.179 (0.974) | -0.09 | 0.956 |
| | | $\gamma_1$ | -1.635 (0.527) | -0.09 | 0.956 |
| | | $\beta_0$ | -1.541 (0.494) | -0.027 | 0.932 |
| | | $\beta_1$ | 1.025 (0.227) | -0.025 | 0.936 |
| | 200 | $\gamma_0$ | 2.091 (0.678) | -0.046 | 0.954 |
| | | $\gamma_1$ | -1.561 (0.354) | -0.041 | 0.968 |
| | | $\beta_0$ | -1.531 (0.322) | -0.021 | 0.916 |
| | | $\beta_1$ | 1.016 (0.15) | -0.016 | 0.914 |

1) Bias (%) = $\frac{\text{true value} - \text{estimated value}}{\text{true value}} \times 100\%$ .
2) CP = $\frac{\sum_{m=1}^{500} I(\text{true value} \in (\delta_{2.5\%}, \delta_{97.5\%}))}{500}$ , $(\delta_{2.5\%}, \delta_{97.5\%})$ is 95% credible interval.

Except *γ*_0_ has 10% bias in Case 2 (*n* = 100), all other biases are less than 5%, and all SDs are small which indicates that our estimation is very stable. All CPs are greater than 90%. Overall, metropolis algorithm performed well on ZIB regression.

## 3. Application to Dose-Finding Study

In this section, we will introduce the application of BZIB regression to dose-finding studies. Assuming we have *N* cohorts. Let *n*_*i*_ denote the number of patients, *y*_*i*_ denote the number of DLTs, and *π*_*i*_ denote the probability of DLT, in *i*th cohort. *p*_*i*_ is the probability that *y*_*i*_ is generated from observation of only zeros. Imposing logit links on *p*_*i*_ and *π*_*i*_, that is, *logit*(*p*_*i*_) = *γ*_0_ + *γ*_1_*dose*_*i*_, and *logit*(*π*_*i*_) = *β*_0_ + *β*_1_*dose*_*i*_. It is reasonable to assume that *p*_*i*_ is decreasing with doses (i.e., the probability of 0 DLT should be getting smaller as the increasement of doses) and *π*_*i*_ is increasing with doses, so we have *γ*_1_ < 0, and *β*_1_ > 0.

### 3.1 Prior Specification

As per our assumption, a below 0 and an above 0 truncated normal prior can be assigned to *γ*_1_ anD *β*_1_, respectively (Tibaldi, 2008). Regular normal priors can be assigned to *γ*_0_ anD *β*_0_, since we have no restrictions for them. Usually, we do not use non-informative prior in dose-finding studies due to small sample size. However, since we lack historical data for this study, by discussing with team, our guesstimates are 1) *p*_1_ is greater than 50%, 2) *p*_8_ should be close to 0, 3) *π*_1_ should be close to 0, 4) *π*_8_ is no less than 50%, 5) 22 mg may be MTD. Based on our guesstimates, the priors we used are: *γ*_0_∼*N*(2.5, 2), *γ*_1_∼*TN*_0_− (−0.1, 2), *β*_0_∼*N*(−5, 2), *β*_1_ ∼*TN*_0_+ (0.1, 0.15). Mean and 95% Credible Interval (CI) of Prior Probabilities of observing only zeros and DLT at each dose are shown in Figure 1. Both curves for *p*_*i*_ and *π*_*i*_ comply with our guesstimates, and the broad 95% CI indicates that our priors are weakly informative.

**Figure 1:**
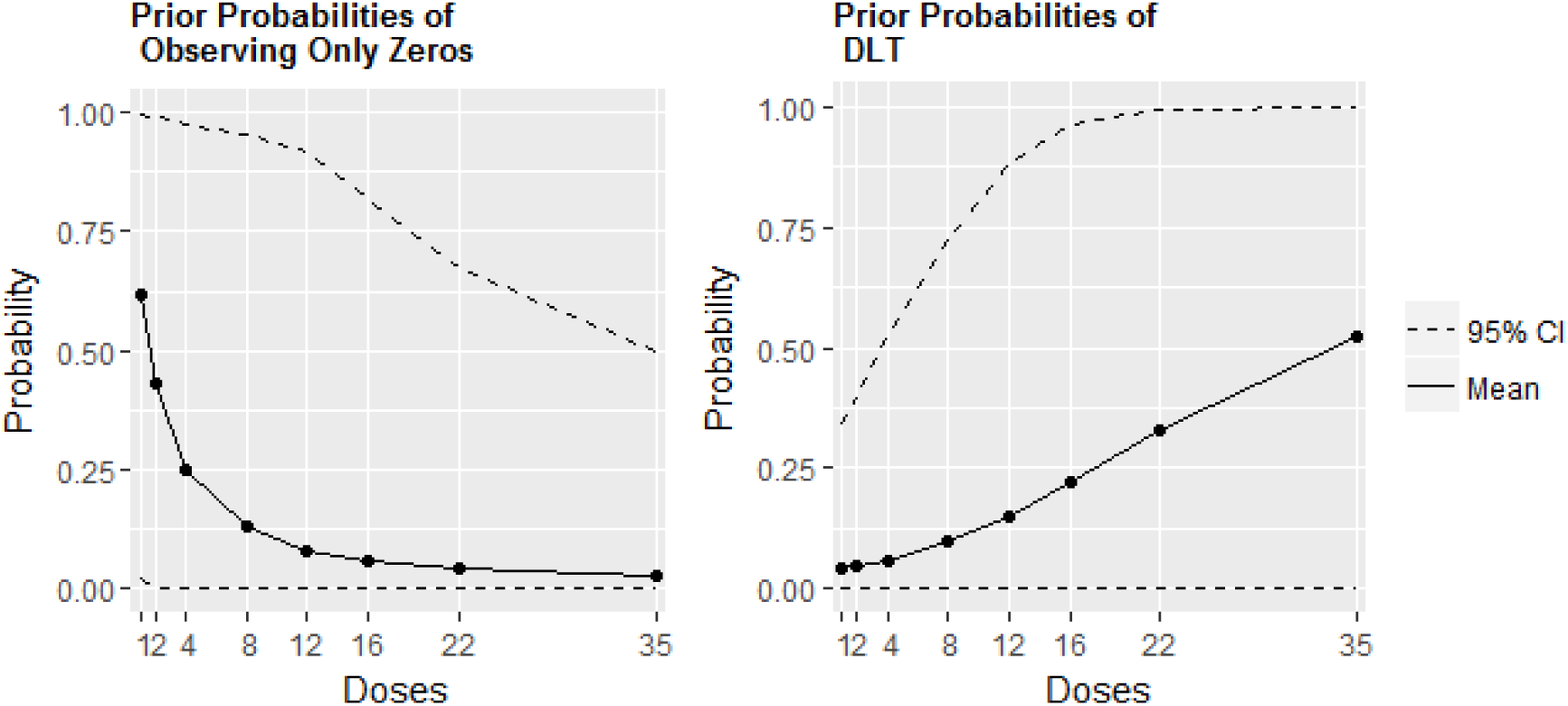
Mean and 95% CI of Prior Probabilities in BZIB regression

### 3.2 Criteria to Select Recommended Dose

We adopted the criteria in Neuenschwander (2008) to select recommended dose based on the MCMC values of *π*_*i*_ sampled from posterior distribution,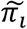 If we categorize estimated probabilities of DLT into three intervals: 1) Under dose interval: (0, 0.16], 2) Target dose interval: (0.16, 0.33], and 3) Over dose interval: (0.33, 1], then the probability of under dose, target dose, and over dose at each dose level will be calculated as:

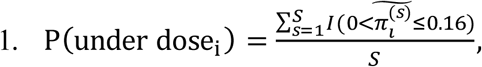

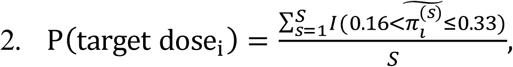

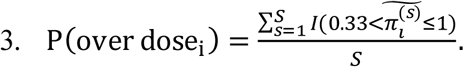

The recommended dose is defined as the dose with P(over dose) ≤ *τ* and maximum P(TargeT dose). *τ* is the threshold of over dose control proposed by Babb (1998). In this paper, we use *τ* = 0.25.

### 3.3 Dose-Finding Simulation

We conducted 1000 dose-finding simulations based on 11 scenarios (S1 – S11) to access the accuracy of our BZIB regression in dose-finding study. We have 8 cohorts in each scenario, and 3 patients will be enrolled each time, and the maximum sample size is 48. Our simulation start enrolling patients from the first cohort and if the recommended dose from BZIB regression is higher than the current dose, we escalate the current dose one more level, otherwise, we deescalate the current dose to recommended dose. When recommended dose is equal to current dose, no less than 6 patients at current dose, and P(target dose) of current dose is greater than or equal to 50%, our simulation stops, and the recommended dose will be claimed as our selected target dose (or MTD in the given doses).

To test the necessity of zero-inflated part in our model, we compared our BZIB regression with two very widely-used logistic regressions in dose-finding studies in pharmaceutical industry, which do not concern the observation of zeros. The description of these two models are below.

#### Model 1

Regular Bayesian logistic regression (RBLR) (Guédé, 2014; Tibaldi, 2008).

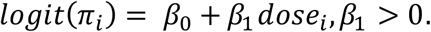

Where, *β*_0_ is imposed with a normal prior, *β*_1_ is imposed with a truncated normal prior.

In our simulation, we gave *β*_0_ and *β*_1_ the same prior as in our BZIB regression.

#### Model 2

Two-parameter Bayesian logistic regression (TBLR) (Neuenschwander, 2008).

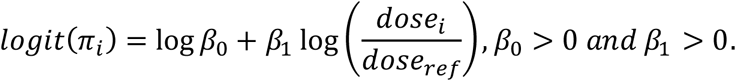

Where, *dose*_*ref*_ is an arbitrary referent dose. (log *β*_0_, log *β*_1_) is imposed with a bivariate normal prior. In our simulation, our *dose*_*ref*_ is 22 mg, and we adopted a weakly informative prior proposed by Neuenschwander (2015), since we lack historical data for this study, and the prior probabilities provided by this prior comply with our guesstimates in Section 3.1. Please see Figure 2.

**Figure 2:**
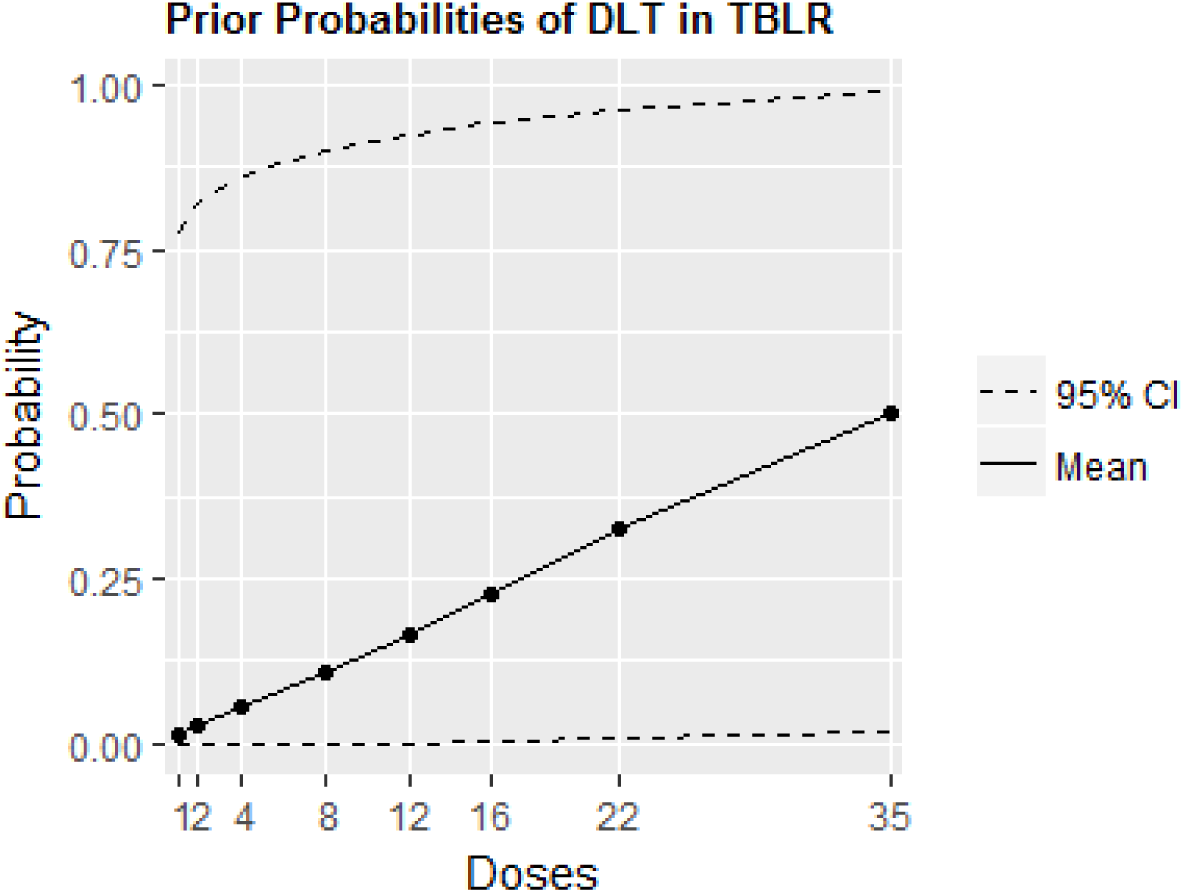
Mean and 95% CI of Prior Probabilities in TBLR.

Table 2 shows the scenarios and the probabilities that observed target doses were selected as MTD with BZIB regression, RBLR, and TBLR. Values for target doses are bold. In Scenario 1, all observed probabilities of DLT are in under dose interval, and all three models selected 35 mg with the highest probability, this is because 35 mg is the highest dose which cannot be escalated. However, RBLR performed very conservatively with just 54.2% at 35 mg. Scenario 2 has no target dose either, and all observed probabilities of DLT are in over dose interval. All three models showed very low probability to select over doses. Scenario 3 and 4 have target doses in high dose part, and no less than half cohorts have no DLT. RBLR was not able to provide adequate accuracy to select target doses in Scenario 4. Scenario 5 has two target doses in the middle part, and our three models provided the similar accuracies. Scenario 6 and 7 have relatively low target doses. In Scenario 6, although BZIB regression has a lower accuracy than RBLR and TBLR, its probability of selecting target dose is still greater than 50%, and furthermore, BZIB regression has lower probability to select over doses than RBLR and TBLR. In Scenario 7, BZIB regression is the only model reached accuracy of 50%. Scenario 8 – 11 have big jumps between target doses and its next doses, and no less than half cohorts have no DLT. In Scenario 8 and 9, BZIB regression has significantly higher accuracy than the other two models. In Scenario 10 and 11, although all models selected target doses successfully, BZIB regression has the highest accuracy.

**Table 2:** Scenarios and Dose-Finding Simulation Results

|  |  | Doses (mg) |  |  |  |  |  |  |  |
| --- | --- | --- | --- | --- | --- | --- | --- | --- | --- |
|  |  | 1 | 2 | 4 | 8 | 12 | 16 | 22 | 35 |
| S1 | Obs. P(DLT) | 0 | 0 | 0.01 | 0.02 | 0.05 | 0.07 | 0.08 | 0.1 |
|  | BZIB Selection (%) | 0.1 | 0.1 | 0.3 | 0.4 | 2.4 | 5.3 | 14.8 | 76.6 |
|  | RBLR Selection (%) | 0 | 0 | 0 | 0.2 | 0.7 | 8.2 | 36.7 | 54.2 |
|  | TBLR Selection (%) | 0 | 0 | 0 | 0.1 | 0.9 | 6.1 | 12 | 80.9 |
| S2 | Obs. P(DLT) | 0.60 | 0.65 | 0.70 | 0.75 | 0.80 | 0.85 | 0.90 | 0.95 |
|  | BZIB Selection (%) | 3.4 | 0.3 | 0.1 | 0 | 0 | 0 | 0 | 0 |
|  | RBLR Selection (%) | 1.1 | 0.8 | 0.1 | 0 | 0 | 0 | 0 | 0 |
|  | TBLR Selection (%) | 0.7 | 0 | 0 | 0 | 0 | 0 | 0 | 0 |
| S3 | Obs. P(DLT) | 0 | 0 | 0 | 0 | 0.12 | **0.27** | 0.43 | 0.56 |
|  | BZIB Selection (%) | 0 | 0 | 0 | 0.4 | 16.5 | **64.8** | 17.8 | 0.5 |
|  | RBLR Selection (%) | 0 | 0 | 0 | 0 | 36 | **59.8** | 4.2 | 0 |
|  | TBLR Selection (%) | 0 | 0 | 0 | 0.1 | 29.3 | **60** | 10.5 | 0.2 |
| S4 | Obs. P(DLT) | 0 | 0 | 0 | 0 | 0 | 0.12 | **0.28** | 0.46 |
|  | BZIB Selection (%) | 0 | 0 | 0 | 0.4 | 0.2 | 33.3 | **60.6** | 5.5 |
|  | RBLR Selection (%) | 0 | 0 | 0 | 0 | 0.2 | 52 | **47.1** | 0.7 |
|  | TBLR Selection (%) | 0 | 0 | 0 | 0 | 0.3 | 44.6 | **52.2** | 2.9 |
| S5 | Obs. P(DLT) | 0.03 | 0.06 | 0.08 | **0.17** | **0.23** | 0.38 | 0.44 | 0.56 |
|  | BZIB Selection (%) | 0 | 0.7 | 9.1 | **34.7** | **41.1** | 12.8 | 1.4 | 0.1 |
|  | RBLR Selection (%) | 0 | 0.6 | 6.5 | **38** | **39.6** | 14.1 | 0.6 | 0 |
|  | TBLR Selection (%) | 0.1 | 1.1 | 11.1 | **32.1** | **35.1** | 16.3 | 3.1 | 0.1 |
|  |  | 1 | 2 | 4 | 8 | 12 | 16 | 22 | 35 |
| S6 | Obs. P(DLT) | 0.03 | 0.07 | **0.18** | 0.39 | 0.45 | 0.53 | 0.61 | 0.7 |
|  | BZIB Selection (%) | 3.9 | 22.3 | **54.8** | 14 | 2.2 | 0.4 | 0 | 0 |
|  | RBLR Selection (%) | 0 | 2.9 | **60.4** | 32.6 | 3.2 | 0.2 | 0 | 0 |
|  | TBLR Selection (%) | 0.1 | 6.2 | **62** | 24.8 | 3.6 | 0.4 | 0 | 0 |
| S7 | Obs. P(DLT) | **0.23** | **0.31** | 0.42 | 0.53 | 0.61 | 0.73 | 0.81 | 0.92 |
|  | BZIB Selection (%) | **34.1** | **21** | 17.3 | 2 | 0 | 0 | 0 | 0 |
|  | RBLR Selection (%) | **7.3** | **23.9** | 21.3 | 1.2 | 0 | 0 | 0 | 0 |
|  | TBLR Selection (%) | **11.2** | **24.2** | 9.6 | 0.8 | 0.1 | 0 | 0 | 0 |
| S8 | Obs. P(DLT) | 0 | 0 | 0 | 0 | 0 | 0.09 | **0.2** | 0.68 |
|  | BZIB Selection (%) | 0 | 0 | 0 | 0.1 | 0.3 | 15.6 | **83.2** | 0.8 |
|  | RBLR Selection (%) | 0 | 0 | 0 | 0 | 0 | 26.9 | **73.1** | 0 |
|  | TBLR Selection (%) | 0 | 0 | 0 | 0 | 0.3 | 25.6 | **73.2** | 0.9 |
| S9 | Obs. P(DLT) | 0 | 0 | 0 | 0 | 0 | 0.11 | **0.27** | 0.89 |
|  | BZIB Selection (%) | 0 | 0 | 0 | 0.7 | 0.7 | 31.3 | **67.2** | 0.1 |
|  | RBLR Selection (%) | 0 | 0 | 0 | 0 | 0.2 | 50 | **49.8** | 0 |
|  | TBLR Selection (%) | 0 | 0 | 0 | 0 | 0.2 | 48.5 | **51.2** | 0.1 |
| S10 | Obs. P(DLT) | 0 | 0 | 0 | 0 | 0.1 | **0.22** | 0.75 | 0.9 |
|  | BZIB Selection (%) | 0 | 0 | 0.2 | 0.7 | 16.5 | **81.4** | 1.2 | 0 |
|  | RBLR Selection (%) | 0 | 0 | 0 | 0 | 25.6 | **74** | 0.4 | 0 |
|  | TBLR Selection (%) | 0 | 0 | 0 | 0.1 | 22.6 | **77** | 0.3 | 0 |
| S11 | Obs. P(DLT) | 0 | 0 | 0 | 0 | 0.09 | **0.18** | 0.84 | 0.95 |
|  | BZIB Selection (%) | 0 | 0 | 0.1 | 0.4 | 12.8 | **85.9** | 0.8 | 0 |
|  | RBLR Selection (%) | 0 | 0 | 0 | 0 | 18.9 | **80.9** | 0.2 | 0 |
|  | TBLR Selection (%) | 0 | 0 | 0 | 0 | 15.8 | **83.9** | 0.3 | 0 |

1) Obs. P(DLT) is observed probability of DLT based on which number of DLTs is generated in each cohort.
2) BZIB Selection, RBLR Selection, and TBLR Selection are the probability of a dose selected as target dose with BZIB regression, RBLR, and TBLR.

All in all, except Scenario 5 and 6, BZIB regression has obviously higher accuracies than RBLR and TBLR. In Scenario 5, all three models have similar accuracies, and in Scenario 6, BZIB regression has better performance in safety control.

### 3.4 Application to an Example

Now, let us apply BZIB regression to the data in our introduction. We ran BZIB regression in R 3.4.3 with 20000 iterations and 10000 burn-ins. Our R code can be found in Appendix A. Our estimates of probabilities of under dose, target dose, over dose, and DLT are shown in Table 3. 25 mg is added as a predicted dose, and assuming 6 patients are enrolled at it. As per our criteria of recommended dose, the P(over dose) of the first 7 doses are less than 0.25, and among them, 22 mg has the maximum P(TargeT dose), so 22 mg is recommended as MTD in our doses. Table 4 shows the estimates of probabilities of *y*∼0 and *y* = 0, we can see that both P(y∼0) and P(*y* = 0) are decreasing with doses, which is in accordance with our assumption. P(*y*_1_∼0) = 0.633, which implies that 0 in the first cohort is more likely from the observation of only zeros than binomial distribution, and all other P(*y*_*i*_∼0)s are small, which indicates that number of DLTs in these cohorts are very likely generated from a binormal distribution. P(y = 0) can be interpreted as the potential possibility that *y* = 0, given *p*_*i*_ and *π*_*i*_. We can see that although 12 mg has no DLT out of 3 patients, it has around 18% of possibility to have at least one DLT.

**Table 3:** Estimates of Probabilities of Under dose, Target dose, Over dose, and DLT

| Doses (mg) | P(under dose) | P(target dose) | P(over dose) | P(DLT) |
| --- | --- | --- | --- | --- |
| 1 | 0.998 | 0.002 | 0 | 0.014 |
| 2 | 0.998 | 0.002 | 0 | 0.016 |
| 4 | 0.996 | 0.004 | 0 | 0.021 |
| 8 | 0.991 | 0.009 | 0 | 0.036 |
| 12 | 0.967 | 0.032 | 0 | 0.064 |
| 16 | 0.806 | 0.19 | 0.004 | 0.112 |
| **22** | **0.186** | **0.609** | **0.205** | **0.252** |
| 25* | 0.035 | 0.418 | 0.548 | 0.356 |
| 35 | 0 | 0.01 | 0.99 | 0.727 |

\*25 mg is a predicted dose.

**Table 4:** Estimates of P(*y*∼0) and P(*y* = 0)

| Doses (mg) | 1 | 2 | 4 | 8 | 12 | 16 | 22 | 25* | 35 |
| --- | --- | --- | --- | --- | --- | --- | --- | --- | --- |
| $P(y_i \sim 0)$ | 0.633 | 0.209 | 0.006 | 0 | 0 | 0 | 0 | 0 | 0 |
| $P(y = 0)$ | 0.985 | 0.962 | 0.939 | 0.895 | 0.821 | 0.489 | 0.175 | 0.071 | 0 |

\*25 mg is a predicted dose.

## 4. Conclusion

In this paper, we provided a very clear Bayesian framework for ZIB regression and its application in DLT-based dose-finding studies. We found that metropolis algorithm performed very stably on BZIB regression via simulations. In our dose-finding simulations, we compared BZIB regression with RBLR and TBLR, and our simulation results show that BZIB regression has better performance when data has excessive zeros, and big jump between target dose and its next dose. Even for the data without excessive zeros, BZIB regression provides higher accuracy in all scenarios with high target doses than the other two models as well, and either better safety control or higher accuracy in scenarios with low target doses.

Additionally, compared with the logistic regressions which do not concern observation of zeros, BZIB regression has more flexibility. First, BZIB regression analyses dose-finding data from two aspects: 1) observation of only zeros, 2) number of DLTs based on binomial distribution, that is, two curves will be fit for data analysis. and when *p* goes to 0, BZIB regression goes to a regular logistic regression. Second, one additional control for selecting recommended dose can be added on P(y = 0) if necessary (i.e., 1 − P(y = 0) ≤ φ, the value of φ should be determined based on the studies).

## Acknowledgements

This work was financially supported by the grant of King Mongkut’sInstitute of Technology Ladkrabang. The authors thank the early phase development team in Celgene Corporation for their help on statistical techniques.

### Appendix A: R code

~~~
#### R package “truncnorm” and “progress” need to be installed
#### library(truncnorm)
library(progress)
~~~

~~~
set.seed(10)
~~~

~~~
#### data ####
n = c(3, 3, 3, 3, 3, 6, 6, 6) # number of patients in each cohort
## n1 is used for prediction ##
n1 = c(3, 3, 3, 3, 3, 6, 6, 6, 6) # assume 6 patients were enrolled at 25 mg
x = c(1, 2, 4, 8, 12, 16, 22, 35) # administered doses
y = c(0, 0, 0, 0, 0, 1, 2, 4) # number of DLTs in each cohort
doses = c(1, 2, 4, 8, 12, 16, 22, 25, 35) # 25 mg is for prediction
~~~

~~~
#### priors ####
### gamma0 ∼ N(2.5, 2) ####
mur0 = 2.5
sigr0 = 2
~~~

~~~
#### gamma1 ∼ TN(−0.1, 2) ####
mur1 = −0.1
sigr1 = 2
~~~

~~~
#### beta0 ∼ N(−5, 2)####
mub0 = −5
sigb0 = 2
~~~

~~~
#### beta1 ∼ TN(0.1, 0.25)####
mub1 = 0.1
sigb1 = 0.15
~~~

~~~
#### number of iteration and burn-in ####
niter = 20000
nburnin = 10000
~~~

~~~
#### log liklihood ####
loglkh = function(x, y, n, r0, r1, b0, b1, u){
 n_fac = factorial(n)
 y_fac = factorial(y)
 ny_fac = factorial(n-y)
 ll =u*log(exp(r0+r1*x)+(1+exp(b0+b1*x))^(-n))-log(1+exp(r0+r1*x))+
  (1-u)*(y*(b0+b1*x)-n*log(1+exp(b0+b1*x))+log(n_fac/(y_fac*ny_fac)))
 return(ll)
}
~~~

~~~
#### sigmoid funcitons ####
sigmoid = function(z){
 return(1/(1+exp(-z)))
}
~~~

~~~
#### initial values ####
r0 = 0
r1 = −0.5
b0 = 0
b1 = 0.5
~~~

~~~
r_0 = r_1 = b_0 = b_1 = NULL
~~~

~~~
#### u ####
u = (y == 0)*1
~~~

~~~
pb <-progress_bar$new(total = niter)
for(i in 1:niter){
 #### updata gamma0 ####
r0_cand = rnorm(1, r0, 1)
logar = sum(loglkh(x, y, n, r0_cand, r1, b0, b1, u))+
 log(dnorm(r0_cand, mur0, sigr0))-
 sum(loglkh(x, y, n, r0, r1, b0, b1, u))-
 log(dnorm(r0, mur0, sigr0))
~~~

~~~
w = runif(1, 0, 1)
~~~

~~~
if(log(w) < logar){r0 = r0_cand}
~~~

~~~
r_0 = c(r_0, r0)
~~~

~~~
#### updata gamma1 ####
r1_cand = rtruncnorm(1, a=-Inf, b = 0, r1, 1)
logar = sum(loglkh(x, y, n, r0, r1_cand, b0, b1, u))+
 log(dtruncnorm(r1_cand, a=-Inf, b=0, mur1, sigr1))-
 sum(loglkh(x, y, n, r0, r1, b0, b1, u))-
 log(dtruncnorm(r1, a=-Inf, b=0, mur1, sigr1))
~~~

~~~
w = runif(1, 0, 1)
~~~

~~~
if(log(w) < logar){r1 = r1_cand }
~~~

~~~
r_1 = c(r_1, r1)
~~~

~~~
#### updata beta0 ####
b0_cand = rnorm(1, b0, 1)
logar = sum(loglkh(x, y, n, r0, r1, b0_cand, b1, u))+
 log(dnorm(b0_cand, mub0, sigb0))-
 sum(loglkh(x, y, n, r0, r1, b0, b1, u))-
 log(dnorm(b0, mub0, sigb0))
~~~

~~~
w = runif(1, 0, 1)
~~~

~~~
if(log(w) < logar){b0 = b0_cand }
~~~

~~~
b_0 = c(b_0, b0)
~~~

~~~
#### updata beta1 ####
b1_cand = rtruncnorm(1, a=0, b=Inf, mean = b1, sd = 1)
logar = sum(loglkh(x, y, n, r0, r1, b0, b1_cand, u))+
 log(dtruncnorm(b1_cand, a = 0, b = Inf, mub1, sigb1))-
 sum(loglkh(x, y, n, r0, r1, b0, b1, u))-
 log(dtruncnorm(b1, a = 0, b = Inf, mub1, sigb1))
~~~

~~~
w = runif(1, 0, 1)
~~~

~~~
if(log(w) < logar){b1 = b1_cand}
~~~

~~~
b_1 = c(b_1, b1)
~~~

~~~
pb$tick()
}
~~~

~~~
#### MCMC values ####
r0_mc = r_0[(nburnin+1):niter]
r1_mc = r_1[(nburnin+1):niter]
b0_mc = b_0[(nburnin+1):niter]
b1_mc = b_1[(nburnin+1):niter]
~~~

~~~
# estimatesof parameters
params = c(mean(r0_mc), mean(r1_mc), mean(b0_mc), mean(b1_mc))
names(params) = c(“gamma0”, “gamma1”,”beta0”, “beta1”)
~~~

~~~
tmp_tox0 = matrix(nrow = length(doses), ncol = niter-nburnin)
tmp_tox = matrix(nrow = length(doses), ncol = niter-nburnin)
pcat = matrix(nrow = length(doses), ncol = 3)
~~~

~~~
for(i in 1:length(doses)){
 tmp_tox[i,] = sigmoid(b0_mc+b1_mc*doses[i])
 pcat[i, 1] = mean(tmp_tox[i, ]<=0.16)
 pcat[i, 2] = mean(tmp_tox[i, ]>0.16 & tmp_tox[i, ]<=0.33)
 pcat[i, 3] = mean(tmp_tox[i, ]>0.33)
}
~~~

~~~
for(i in 1:length(doses)){
 tmp_tox0[i, ] = sigmoid(r0_mc+r1_mc*doses[i])
}
~~~

~~~
ptox = apply(tmp_tox, 1, mean)
result = round(cbind(doses, pcat, ptox), 3)
colnames(result) = c(“doses”, “punder”, “ptarget”, “pover”, “pdlt”)
~~~

~~~
#### P(y∼0) and P(y=0) ####
py_0 = round(sigmoid(params[1]+params[2]*doses), 3)
pye0 = round(py_0 + (1-py_0)*(1-ptox)^n1, 3)
py0 = rbind(py_0, pye0)
colnames(py0) = doses
rownames(py0) = c(“P(y∼0)”, “P(y=0)”)
~~~

~~~
#### plot MCMC values ####
par(mfrow = c(2, 2))
plot(r0_mc, type=“l”)
plot(r1_mc, type=“l”)
plot(b0_mc, type=“l”)
plot(b1_mc, type=“l”)
~~~

~~~
#### summary ####
summary = list(“summary of P(DLT)” = round(result, 4), “P(y∼0) and P(y=0)” = py0, “Estimates of Parameters” = params)
summary
~~~

